# Expression of TorsinA in a heterologous yeast system reveals interactions with conserved lumenal domains of LINC and nuclear pore complexes

**DOI:** 10.1101/421909

**Authors:** Madeleine Chalfant, Karl W. Barber, Sapan Borah, David Thaller, C. Patrick Lusk

## Abstract

DYT1 dystonia is caused by an in-frame deletion of a glutamic acid codon in the gene encoding the AAA+ ATPase TorsinA. TorsinA localizes within the lumen of the nuclear envelope/ER and binds to a membrane-spanning co-factor, LAP1 or LULL1, to form an ATPase; the substrate(s) of TorsinA remain ill defined. Here we use budding yeast, which lack Torsins, to interrogate TorsinA function. We show that TorsinA accumulates at nuclear envelope embedded spindle pole bodies (SPBs) in a way that requires its oligomerization and the conserved SUN-domain protein, Mps3. TorsinA is released from SPBs upon expression of LAP1 and stabilized by LAP1 mutants incapable of stimulating TorsinA ATPase activity, suggesting the recapitulation of a TorsinA-substrate cycle. While the expression of TorsinA or TorsinA-ΔE impacts the fitness of strains expressing *mps3* alleles, a genetic interaction with a conserved component of the nuclear pore complex, Pom152, is specific for TorsinA. This specificity is mirrored by a physical interaction between Pom152 and TorsinA, but not TorsinA-ΔE. These data suggest that TorsinA-nucleoporin interactions would be abrogated by TorsinA-ΔE, providing new experimental avenues to interrogate the molecular basis behind nuclear envelope herniations seen in cells lacking TorsinA function.

## INTRODUCTION

DYT1 dystonia is an early-onset, heritable movement disorder caused by an autosomal dominant mutation removing a glutamic acid codon (ΔE) in the *DYT1/TOR1A* gene that encodes the AAA+ ATPase TorsinA (TorA)(Ozelius *et al.*, 1997). How expression of the TorsinA-ΔE/TorA-ΔE allele, and several others (Ozelius *et al.*, 1997; Zirn *et al.*, 2008; Xiao *et al.*, 2009; Rebelo *et al.*, 2015), causes dystonia remains ill defined and requires additional molecular insight into the function of TorA. It remains unclear whether TorA is a typical AAA+ ATPase, which act as oligomeric ring assemblies that perform mechanical work on protein or protein complex substrates (Hanson and Whiteheart, 2005) as there is no clear consensus as to the identity of TorA substrates.

Challenges with identifying TorA substrates likely relate to the fact that TorA is an unusual member of the AAA+ family. For example, it is uniquely localized within the contiguous lumen of the nuclear envelope (NE) – endoplasmic reticulum (ER) system (Hewett *et al.*, 2000; Kustedjo *et al.*, 2000). Further, while TorA has canonical AAA+ ATPase motifs (Ozelius *et al.*, 1997), its ATP binding pocket lacks the arginine finger (Brown *et al.*, 2014; Sosa *et al.*, 2014) required for ATP hydrolysis (Ogura *et al.*, 2004). To form a functional enzyme, TorA must bind to one of two membrane-spanning co-factors, LAP1 or LULL1, that contribute this critical arginine finger to TorA’s ATP binding pocket (Zhao *et al.*, 2013; Brown *et al.*, 2014; Sosa *et al.*, 2014); TorA-ΔE is unable to bind to LAP1 or LULL1 suggesting that dystonia is caused by a loss of ATPase function. (Naismith *et al.*, 2009; Zhu *et al.*, 2010; Zhao *et al.*, 2013; Demircioglu *et al.*, 2016). Of note, LAP1 and LULL1 have distinct spatial distributions (Goodchild and Dauer, 2005), with LULL1 having access to the entire ER (Goodchild *et al.*, 2015) and LAP1 being restricted to the inner nuclear membrane (INM)(Senior and Gerace, 1988). This suggests the compelling possibility that these co-factors may influence TorA ATPase activity (or potential substrates) in distinct subdomains of the ER, including the NE.

The idea that TorA has a NE-specific role is bolstered by the appearance of NE herniations or blebs found in developing neurons of TorA^-/-^ and TorA^ΔE/ΔE^ mice (Goodchild *et al.*, 2005; Tanabe *et al.*, 2016). In most cases, these NE herniations, which have been visualized in several other model systems where TorA function is perturbed, bloom from an electron-dense structure at the INM of enigmatic origin (Naismith *et al.*, 2004; Kim *et al.*, 2010; Jokhi *et al.*, 2013; VanGompel *et al.*, 2015; Laudermilch *et al.*, 2016; Tanabe *et al.*, 2016). Recently, electron tomography of HeLa cells lacking all four Torsin isoforms reveals that the electron-dense structures are morphologically similar to nuclear pore complexes (NPCs) and can be labeled with anti-nucleoporin antibodies (Laudermilch *et al.*, 2016). As similar herniations form over defective NPCs in budding yeast in response to perturbations in NPC assembly and/or the triggering of NPC quality control pathways (Wente and Blobel, 1993, 1994; Aitchison *et al.*, 1995; Murphy *et al.*, 1996; Siniossoglou *et al.*, 1996; Zabel *et al.*, 1996; Emtage *et al.*, 1997; Ryan and Wente, 2002; Webster *et al.*, 2014; Webster *et al.*, 2016; Onischenko *et al.*, 2017; Thaller and Lusk, 2018; Zhang *et al.*, 2018), it seems reasonable that TorA might play a role in the biogenesis of NPCs. It remains unclear, however, what components of the NPC could be targeted by TorA, although gp210/Nup210 is an obvious candidate given its massive (Wozniak *et al.*, 1994) and well conserved lumenal domain (Upla *et al.*, 2017).

It is equally likely, however, that TorA might indirectly influence NPC assembly by, for example, altering the function of proteins that contribute to lipid metabolism (Grillet *et al.*, 2016) or by binding and modulating other NE components like LINC (Linker of Nucleoskeleton and Cytoskeleton) complexes, which have been implicated in NPC biogenesis from yeast to man (Liu *et al.*, 2007; Lu *et al.*, 2008; Friederichs *et al.*, 2011; Talamas and Hetzer, 2011; Chen *et al.*, 2014). LINC complexes are composed of trimers of INM SUN (Sad1 and UNc-84) and outer nuclear membrane (ONM) KASH (Klarsicht, Anc-1, and Syne homology)-domain containing proteins that interact within and span the perinuclear space, physically coupling the cytoskeleton and nucleoskeleton (Padmakumar *et al.*, 2005; Crisp *et al.*, 2006; Sosa *et al.*, 2012). Indeed, the concept that TorA might remodel LINC complexes remains a compelling narrative, supported by several biochemical, genetic and cell biological data (Nery *et al.*, 2008; Vander Heyden *et al.*, 2009; Jungwirth *et al.*, 2011; Atai *et al.*, 2012; Saunders *et al.*, 2017; Dominguez Gonzalez *et al.*, 2018).

Here, we take advantage of a heterologous budding yeast genetic system to interrogate the function of TorA. While yeast lack a TorA orthologue, they express several evolutionary conserved NE components including NPCs, LEM domain integral INM proteins, and at least one SUN domain protein, Mps3 (Webster and Lusk, 2016). Our data are consistent with a model in which human TorA can interact with and impact the function of yeast NE proteins with conserved lumenal domains, including Mps3.Moreover, both genetic and physical interaction data support that TorA can interact with the Nup210 orthologue, Pom152. That this interaction is lost with TorA-ΔE point to a new perspective on how to interpret the underlying causes of the NE abnormalities associated with early onset dystonia.

## RESULTS AND DISCUSSION

### TorA accumulation at SPBs requires oligomerization

Several prior studies have expressed TorA in budding yeast, but no clear functional insight for TorA has emerged (Valastyan and Lindquist, 2011; Zacchi *et al.*, 2014; Adam *et al.*, 2017; Zacchi *et al.*, 2017). However, given that these studies were largely performed before the discovery of the LAP1 and LULL1 co-factors required for TorA ATPase activity, we felt it was worthwhile to revisit this system using the fully functional enzyme complex. We took advantage of TorA-GFP constructs generated by Valastyan et al (2011), which were engineered with a yeast-specific promoter and a Kar2 signal sequence (Figure 1A). To reduce variation in cell-to-cell expression levels, we modified these constructs to allow for chromosomal integration and generated yeast strains expressing TorA-GFP and several TorA alleles including TorA-ΔE-GFP, TorA-EQ-GFP and TorA-GD-GFP (Figure 1A).

**Figure 1.**
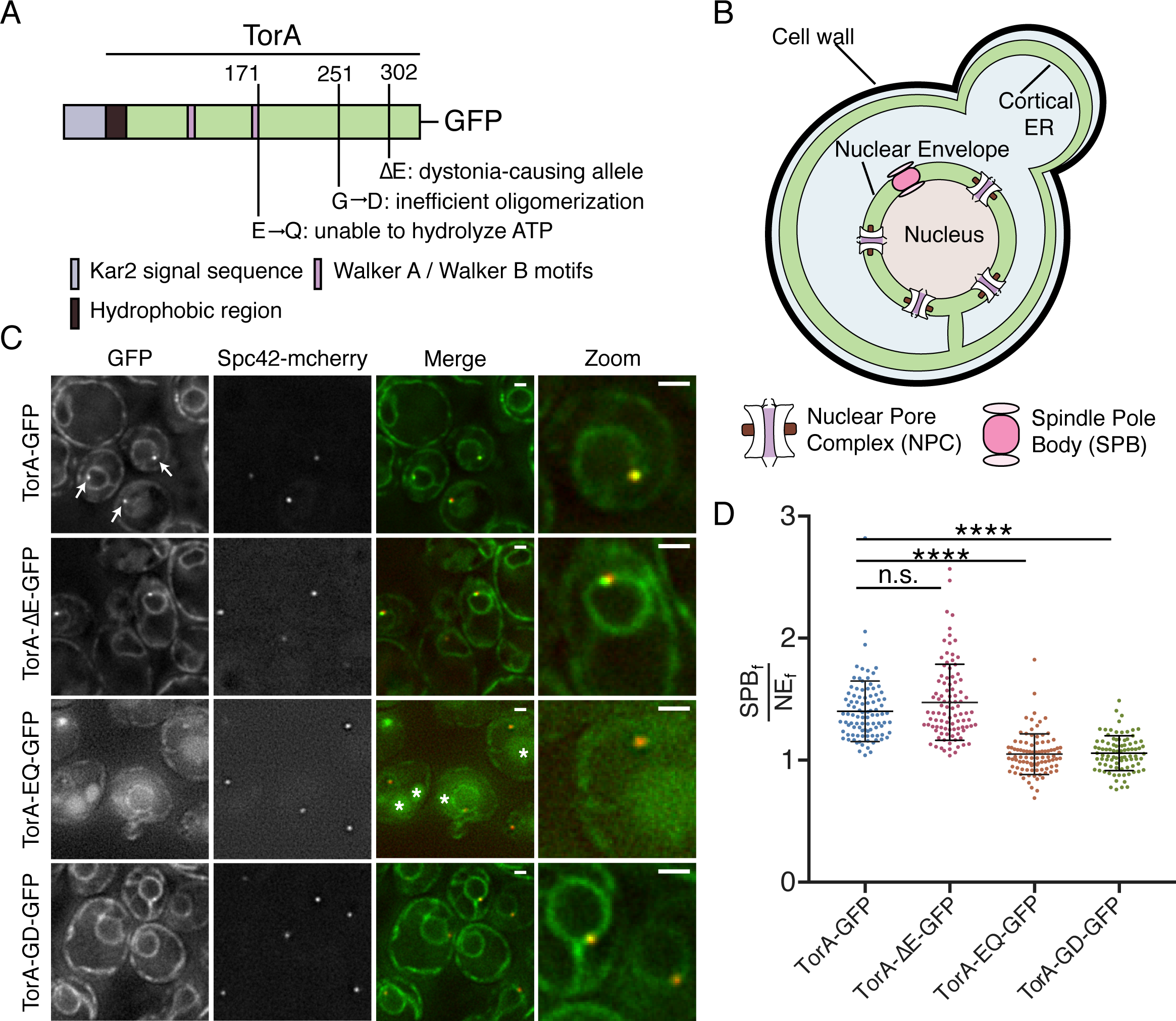
TorA-GFP and TorAΔE-GFP, but not TorA-GD-GFP, accumulate at SPBs. (A) Schematic of TorA-GFP constructs - relevant motifs and locations (numbers are amino acid residue positions from the start methionine of the human gene) of amino acid changes are shown. (B) Diagram (not to scale) of a budding yeast cell schematizing the NE/ER system with NPCs and SPBs. (C) Deconvolved fluorescent micrographs of strains expressing the indicated TorA-GFP constructs and Spc42-mCherry. Green and red fluorescent images alongside a merge and magnification of one cell (zoom) are shown. Arrows point to NE foci that colocalize with Spc42-mCherry. TA-EQ-GFP might accumulate in vacuoles (asterisks). Bar is 1 µm. (D) Plot of SPB_f_/NE_f_ of indicated TA-GFP constructs in individual cells from three biological replicates (n=32/replicate) with mean and SD. Kruskal Wallace ANOVA with *post hoc* Dunn’s test. ****=p<0.0001.

We first examined the localization of TorA-GFP in logarithmically growing wildtype (wt) cells. As previously published (Valastyan and Lindquist, 2011), TorA-GFP was found in a perinuclear (i.e. nuclear envelope) and cortical distribution, consistent with the morphology of the budding yeast ER (Figure 1B). We remarked, however, that TorA-GFP also accumulated in one or two foci at the NE (Figure 1C, arrows). This localization was particularly striking in cells expressing low levels of TorA-GFP, best observed upon growing cells to saturation (Figure S1A). In these cases, we observed TorA-GFP in one or two puncta per cell with a nearly undetectable pool in the rest of the NE/ER, raising the possibility that TorA preferentially binds to a NE-specific structure.

The focal accumulation of TorA-GFP at the NE was reminiscent of spindle pole bodies (SPBs), the yeast centrosome equivalents that span both membranes of the NE (Jaspersen and Ghosh, 2012)(Figure 1B). To test this idea, we examined the localization of TorA-GFP in a strain expressing a mCherry-tagged core component of the SPB, Spc42. As shown in Figure 1C, we observed clear coincidence between virtually all Spc42-mCherry and TorA-GFP NE-foci, confirming that TorA-GFP likely associates with SPBs. In these logarithmically growing cells, we also compared the mean fluorescence of TorA-GFP at the SPB (SPB_f_) with the broader NE (NE_f_) on an individual cell basis to provide a metric of relative SPB enrichment (SPB_f_/NE_f_), which ranged from 1.04 to 2.82 and had an average SPB_f_/NE_f_ of 1.40 (Figure 1D).

We next tested whether TorA-ΔE, TorA-EQ and TorA-GD would also enrich at SPBs. While TorA-ΔE-GFP was produced at lower levels than TorA-GFP (Figure S1B), it nonetheless accumulated at SPBs (mean SPB_f_/NE_f_ of 1.47), similar to its wt counterpart. In contrast, TorA-EQ-GFP did not enrich at SPBs (mean SPB_f_/NE_f_ =1.05) although we struggled to find conditions in which TorA-EQ-GFP was stably expressed - note that even the NE/ER signal was low and there was green fluorescence in the vacuole (see asterisks and Figure S1A) that could indicate its degradation. Interestingly, in the absence of LAP1 or LULL1, TorA-EQ can aggregate *in vitro* (Sosa *et al.*, 2014), which may explain its potential targeting for degradation in our system. Strikingly, however, a TorA-GD mutant, which cannot oligomerize due to disruption of the “back interface” (Chase *et al.*, 2017), failed to accumulate at SPBs (mean SPB_f_/NE_f_=1.06) despite being expressed at levels similar to TorA-ΔE and TorA-EQ (Figure S1B). This was particularly obvious when comparing these strains grown to saturation (Figure S1A). Taken together, these results suggest that TorA might interact with a SPB component in a manner that requires its oligomerization.

### LAP1-LD expression releases TorA from SPBs

There is a general consensus that TorA can self-assemble into an oligomer in its ATP-bound form (Vander Heyden *et al.*, 2009; Jungwirth *et al.*, 2010; Chase *et al.*, 2017).Interestingly, recent work also indicates that binding of the LAP1 lumenal domain (LAP1-LD) to a TorA hexamer and subsequent ATP hydrolysis causes oligomer disassembly (Chase *et al.*, 2017). We took advantage of these observations to attempt to recapitulate a putative ATPase cycle *in vivo*, using TorA-GFP SPB accumulation as a proxy for a substrate interaction. While efforts to produce full length LAP1 in yeast were unsuccessful, by replacing the N-terminus of LAP1 with a fragment of ubiquitin (NubG), we could express the NubG-LAP1-LD (indicated as LAP1-LD, below) under the control of a galactose-inducible (*GAL1*) promoter (Figure 2A), which reached peak levels after ∼5 hours of growth in the presence of galactose (Figure S1C).

**Figure 2.**
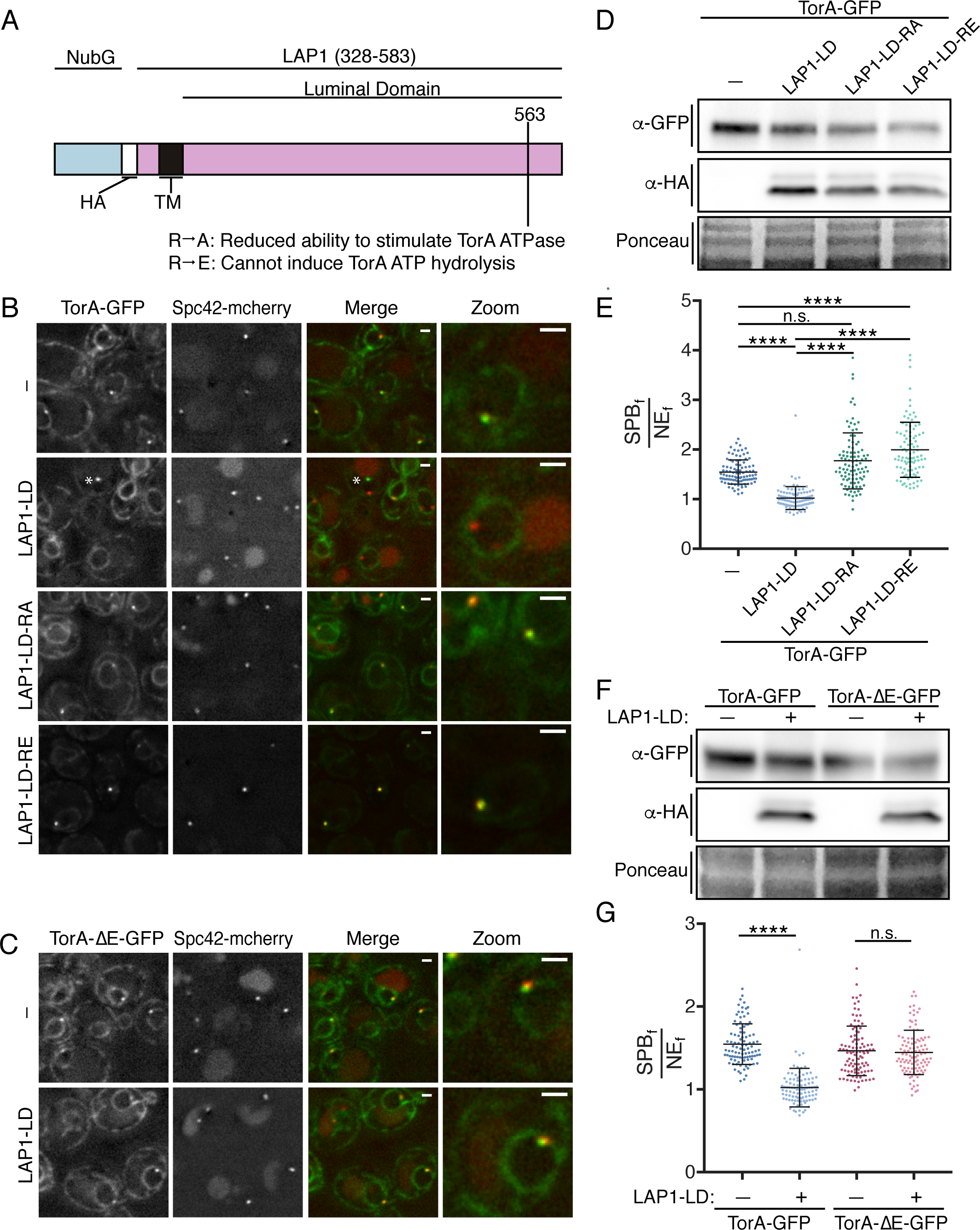
LAP1-LD mediated release of TorA-GFP from SPBs likely requires ATP hydrolysis. (A) Diagram of the LAP1-LD constructs with N-terminal NubG fragment, HA epitope, transmembrane (TM) segment, and position of indicated mutations. (B) Deconvolved fluorescent micrographs of strains expressing TorA-GFP and Spc42-mCherry with or without (-) the production of LAP1-LD constructs. Green and red fluorescence alongside a merge and magnification of a cell (zoom) are shown. Asterisk points to a rare NE focus that is not a SPB only seen upon LAP1-LD expression. (C) As in B with TorA-ΔE-GFP. Bar is 1 µm (D) Western blot of TorA-GFP (α-GFP) and LAP1-LD constructs (α-HA) with Ponceau stain to assess loads. (E) Plot of SPB_f_/NE_f_ of experiment in (B) of individual cells from three biological replicates (n=32/replicate) with mean (middle line) and SD. Kruskal Wallace ANOVA with *post hoc* Dunn’s test. ****=p<0.0001. (F) Western blot of TorA/ΔE-GFP (α-GFP) and LAP1-LD (α-HA) levels with reference to Ponceau stain of total protein loads.(G) As in E with indicated expression constructs.

Remarkably, at the highest levels of LAP1-LD expression, TorA-GFP was no longer visibly concentrated at SPBs (Figure 2B and E; mean SPB_f_/NE_f_=1.02). Importantly, this reduction in SPB_f_/NE_f_ values was due to lower SPB_f_ and not higher NE_f_ signal, as TorA-GFP levels remained unaltered upon production of the LAP1-LD (Figure 2D). Further, and consistent with the idea that the ability of LAP1-LD to reduce TorA association with the SPB is direct, LAP1-LD expression at similar levels (Figure 2F) had no effect on the SPB-accumulation of TorA-ΔE-GFP, which is unable to interact with the LAP1-LD (Naismith *et al.*, 2009; Zhu *et al.*, 2010; Zhao *et al.*, 2013)(Figure 2C and G).

LAP1 acts as a co-factor for TorA by contributing a critical arginine residue (at position 563) to the TorA-ATP binding pocket (Brown *et al.*, 2014; Sosa *et al.*, 2014; Demircioglu *et al.*, 2016) changing this “arginine finger” to either alanine or glutamic acid reduces or abolishes ATP hydrolysis, respectively (Brown *et al.*, 2014)(Figure 2A). We therefore tested if the ability of LAP1 to stimulate TorA ATP hydrolysis is required to prevent SPB accumulation of TorA-GFP. In striking contrast to LAP1-LD, at equivalent expression levels (Figure 2D) LAP1-LD-R563A and, in particular, LAP1-LD-R563E not only failed to negatively affect SPB accumulation of TorA-GFP, but instead resulted in higher TorA-GFP accumulation at the SPB (SPB_f_/NE_f_ values reached 3.90, mean of 2.00, Figure 2B, E). These data suggest that the introduction of the LAP1-LD mutants lead to the “trapping” of a potential TorA substrate. In addition, as the SPB_f_/NE_f_ values of TorA-GFP in the LAP1-LD mutants exceed those of TorA-GFP when expressed alone, these data raise the intriguing possibility that there is a co-factor endogenous to yeast capable of stimulating ATP hydrolysis. Such a model predicts that inhibition of this putative co-factor would mirror the hyperaccumulation of TorA at SPBs observed upon expressing the LAP1-R563A/E alleles. Regardless, in the aggregate, these data support that TorA-GFP is capable of undergoing an ATP-hydrolysis cycle by binding and releasing a substrate that is a likely component of the SPB.

### TorA and TorA-ΔE specifically interact with NE proteins

The ability of the “back interface” mutation to disrupt accumulation of TorA-GFP at the SPB and the release of TorA-GFP from the SPB upon expression of LAP1-LD suggest that TorA may bind to a substrate at the SPB. To facilitate the identification of potential TorA binding partners, we affinity purified TorA-GFP and TorA-ΔE-GFP from cryo-lysates derived from wt cells using anti-GFP nanobody-coupled magnetic beads. As shown in Figure 3A, we pulled out both TorA-GFP and TorA-ΔE-GFP alongside at least one major binding partner of ∼70 kD, which we identified by mass spectrometry (MS) to be Kar2 (the orthologue of the ER chaperone, BiP). This result is consistent with previous work and suggests Kar2 plays a critical role in ensuring ER translocation and/or stability of these human proteins (Zacchi *et al.*, 2014). It also remains formally possible that TorA forms a functional complex with Kar2 and contributes to protein quality control pathways in the ER.

**Figure 3:**
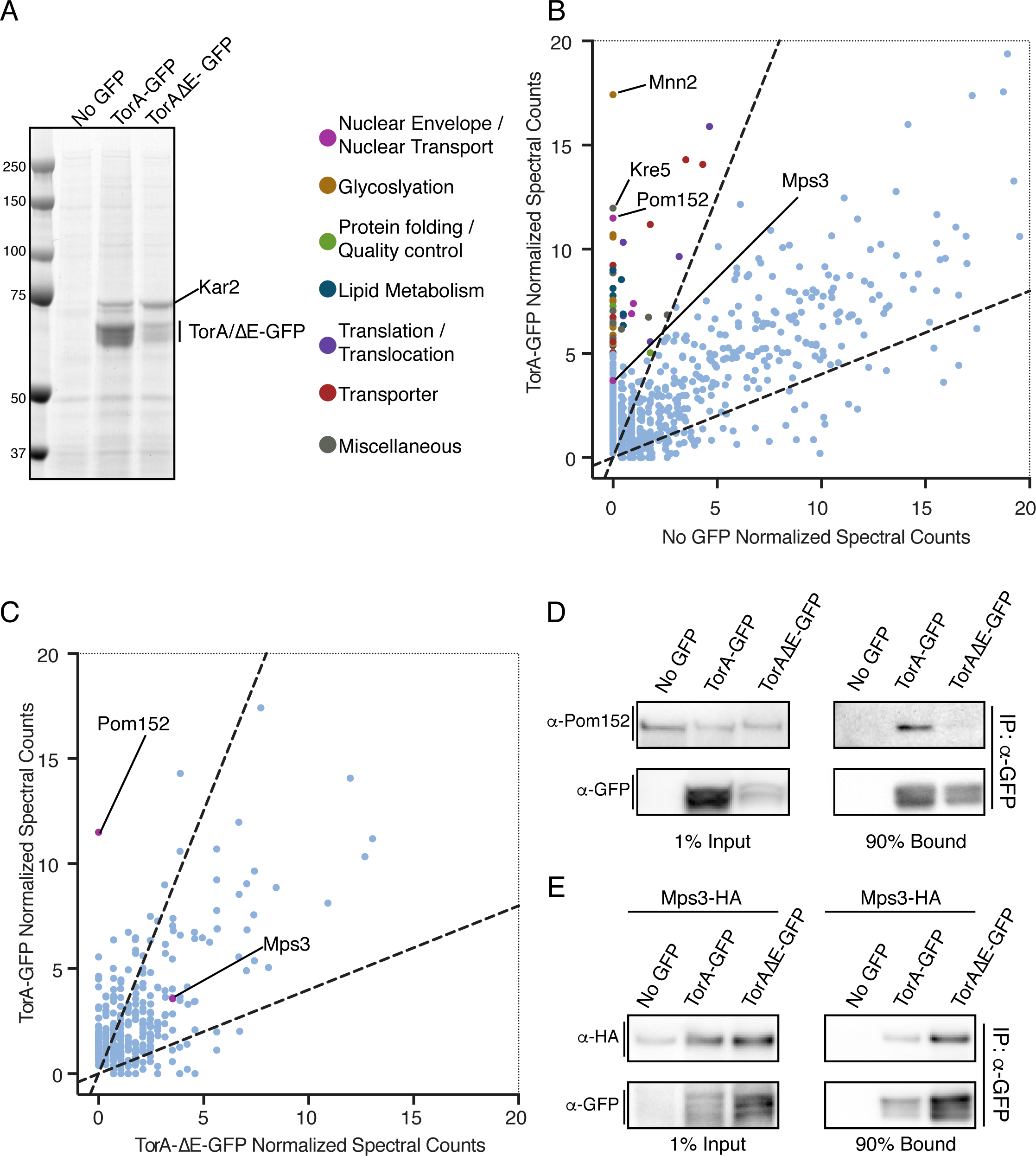
TorA and TorA-ΔE interact with a shared set of proteins but TorA specifically binds to Pom152. (A) Coomassie-stained SDS-PAGE gel of bound proteins eluted from α-GFP nanobody beads incubated with cell extracts expressing TorA-GFP, TorA-ΔE-GFP or “no GFP”. Position of MW standards on left. (B) Magnification of plot (full plot in Figure S2A) comparing average normalized MS spectra from 3 replicates identified in TorA-GFP versus No GFP control. Dashed lines demarcate 2.5-fold enrichment. Proteins with an average of 5 or more spectra that were 2.5-fold enriched in TorA-GFP sample or absent in no GFP sample are colored based on the classifications indicated.(C) As in B (full plot in Figure S2B) but comparing average normalized spectra (from 2 replicates) between TorA-GFP and TorA-ΔE-GFP affinity purifications. (D, E) Western blots of input and bound fractions from affinity purifications of TorA-GFP, TAΔE-GFP with no GFP control.

Next, to fully identify the TorA interactome, we subjected the entire TorA-GFP and TorA-ΔE-GFP bound fractions to shotgun LC-MS/MS. As expected from these highly sensitive approaches, we identified hundreds of proteins, most of which were non-specific and present in the “no GFP” control (complete peptide lists can be found in Supplemental Table 1, “all normalized spectra” tab). We therefore plotted mean normalized spectral counts identified from three independent affinity purifications and considered proteins identified from peptides that were at least 2.5 fold enriched in the TorA-GFP bound fractions relative to the no GFP control (Figure S2A, dotted lines) to be likely interactors. To facilitate visualizing proteins less abundant than Kar2, we have magnified the region of the plot with the majority of specific proteins (Figure S2A, box) and have color-coded those with an average of more than 5 peptides based on a manual functional classification scheme (Figure 3B). Strikingly, while the top two hits identified in the no GFP control specific to TorA-GFP were Mnn2 and Kre5 (two highly abundant enzymes involved in glycosylation), the third protein with the most peptides (with none found in no GFP control) was Pom152 (Figure 3B). Pom152 is the orthologue of Nup210 and is the only component of the NPC with a large (>10 kD) lumenal domain. We could also detect at least one additional nucleoporin, Nup192, and two nuclear transport factors, but these had fewer peptides with some in the no GFP control (Supplementary Table 1, TorA vs. no GFP tab). Remarkably, the only other lumenal-domain-containing NE protein identified in all replicates of TorA-GFP pull outs was the SUN-domain containing protein, Mps3. Critically, Mps3 is the only component of the SPB identified in any replicate of TorA-GFP purifications, strongly supporting the specificity of this interaction.

To identify putative factors that might specifically interact with TorA-GFP but not with TorA-ΔE-GFP, we directly compared normalized spectral counts from two independent TorA-GFP and TorA-ΔE-GFP affinity purifications (Figure 3C and S2C). Here, Pom152 immediately stood out. In fact, we did not identify any peptides for Pom152 in either replicate of TorA-ΔE-GFP (Figure S2B, Supplementary Table 1, TorA-ΔE vs. no GFP tab), a result that we further confirmed by Western blotting using anti-Pom152 antibodies of additional affinity purifications (Figure 3D). While there were several other proteins that also show specificity to TorA-GFP (and a few to TorA-ΔE-GFP as well, see Supplemental Table 1), fewer peptides for each were identified and the potential functional relevance of these proteins with respect to TorA or TorA-ΔE are not obvious and difficult to ascribe. By contrast, peptides specific for Mps3 were also identified in both replicates of the TorA-ΔE-GFP pullouts and through additional affinity purifications from cell extracts derived from strains expressing Mps3-HA (Figure 3E), enabling Mps3 detection by Western blotting with anti-HA antibodies. Thus, TorA and TorA-ΔE interact with Mps3, but only TorA specifically binds to Pom152.

### TorA/TorA-ΔE specifically impact the fitness of *mps3* and *pom152* strains

The localization of TorA in an oligomerization and ATPase activity dependent manner to the SPB combined with the identification of Pom152 and Mps3 as likely TorA binding partners raises the possibility that TorA could influence the function of these NE proteins. We therefore tested whether TorA expression impacted the fitness of yeast strains with alleles of *POM152* and *MPS3*. For these experiments, we placed TorA and TorA-ΔE under the control of the conditional *GAL1* promoter as we observed progressive loss of TorA-GFP expression upon serial culturing in some strain backgrounds. Consistent with our hypothesis and biochemistry, strains null for *POM152* were specifically sensitive to the expression of TorA but not TorA-ΔE (Figure 4A). This result suggests that TorA acts as a dominant negative in the absence of Pom152, perhaps by binding and inhibiting an essential factor that lacks the capacity to interact with TorA-ΔE. Alternatively, particularly in light of our biochemical analysis, we favor a model where Pom152 could directly interact with TorA in a way that modulates its activity.

**Figure 4:**
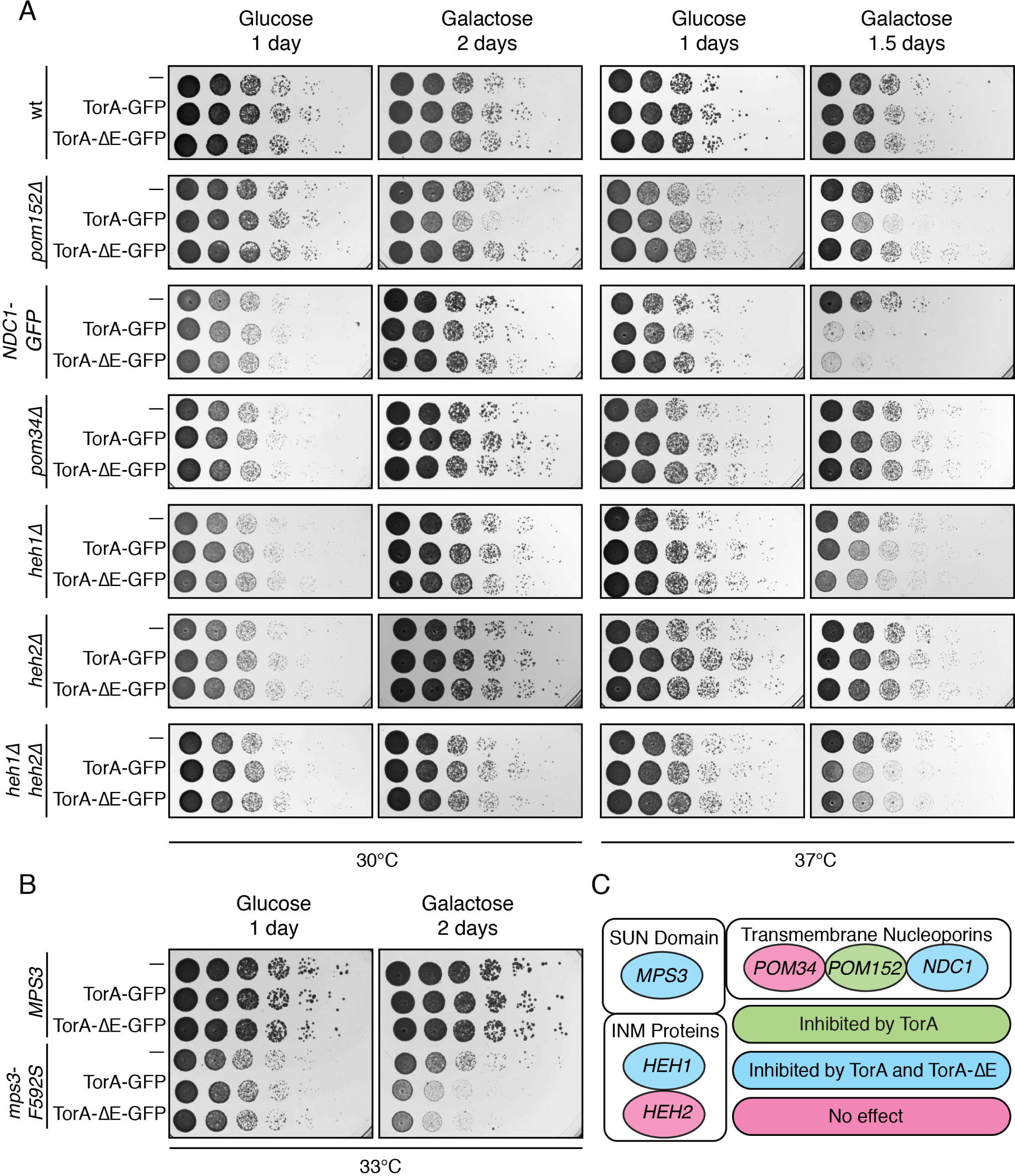
Conditional expression of TorA-GFP or TorAΔE-GFP differentially impacts the fitness of *mps3* and *pom152* strains. (A, B) The indicated strains were 10-fold serially diluted onto glucose or galactose-containing plates, which repress or induce the expression of TorA/TorA-ΔE, respectively. Colony size was assessed after growth at indicated temperatures/time. (C) Summary of genetic backgrounds inhibited by TorA/ TorAΔE expression.

We next tested whether conditional expression of TorA impacted the growth of a *mps3Δ* strain complemented by either *MPS3* or a temperature sensitive allele (*mps3-F592S)* that disrupts the SUN domain and the function of Mps3 (Jaspersen *et al.*, 2006).Interestingly, while expression of TorA or TorA-ΔE did not alter growth of *MPS3*- containing cells, both specifically impacted the fitness of strains expressing the *mps3-F592S* allele (Figure 4B), suggesting that TorA can further perturb a compromised SPB and/or impaired SPB insertion mechanism. As Mps3 likely contributes to both SPB insertion and NPC biogenesis into the NE (Jaspersen *et al.*, 2002; Jaspersen *et al.*, 2006; Friederichs *et al.*, 2011; Jaspersen and Ghosh, 2012; Chen *et al.*, 2014), we tested whether shared components of the SPB and NPC (e.g. Ndc1)(Winey *et al.*, 1993; Chial *et al.*, 1998) are also impacted by TorA expression. For this experiment, we took advantage of our prior observation that C-terminal tagging of Ndc1 with GFP partially impairs its function (Yewdell *et al.*, 2011) and, consistent with this, TorA and TorA-ΔE inhibited the growth of Ndc1-GFP expressing cells, particularly at 37ºC (Figure 4A). Thus, while TorA also genetically interacts with Mps3, in this case this interaction is not affected by the TorA-ΔE allele, again consistent with our biochemical analysis (Figure 4C).

### *mps3* and *pom152* alleles differentially affect SPB accumulation of TorA

Lastly, we tested whether disruption of Mps3 and Pom152 function altered the distribution of TorA-GFP at the SPB. Interestingly, we observed lower SPB_f_/NE_f_ ratios of both TorA-GFP and TorA-ΔE-GFP at the SPB in *mps3-F592S* cells compared to the wt *MPS3* counterpart, even at permissive growth temperatures (RT)(Figure 5A, B and F). These data open up the possibility that TorA (and TorA-ΔE) is recruited to the SPB through a direct interaction with the Mps3 SUN domain. In contrast to the loss of SPB recruitment observed in *mps3-F592S* cells, we observed higher SPB_f_/NE_f_ values for TorA-GFP (up to 3.53, mean SPB_f_/NE_f_ of 2.05) in *pom152Δ* cells (Figure 5C and H), which resembles the hyperaccumulation of TorA-GFP at SPBs seen upon expression of the LAP1-LDs deficient in stimulating TorA-GFP ATP hydrolysis (Figure 2B). Perhaps most interestingly, this effect was specific, and we observed no change in SPB_f_/NE_f_ levels of TorA-ΔE-GFP in *pom152Δ* cells (Figure 5D and H). These data thus closely mirror the genetic and biochemical analysis and support the concept that Pom152 could impact TorA function analogously to its established activator, LAP1.

**Figure 5:**
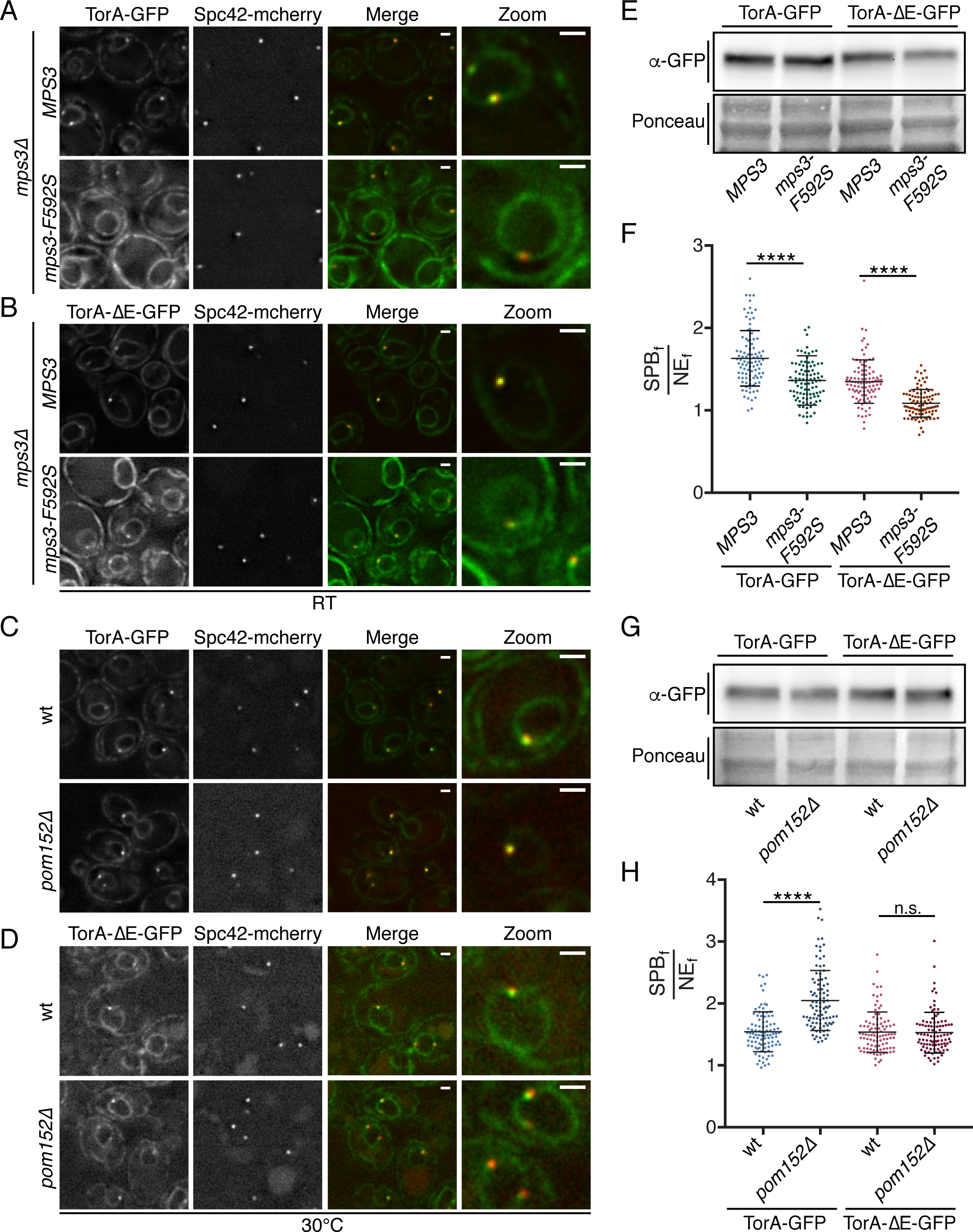
TorA and TorA-ΔE do not enrich at SPBs in *mps3* strains while TorA specifically hyperaccumulates in cells lacking *POM152*. (A-D) Deconvolved fluorescent micrographs of cells expressing TorA-GFP or TorA-ΔE-GFP with Spc42-mCherry in the indicated genetic backgrounds. Green and red fluorescence alongside a merge and magnification of one cell (zoom) are shown. Bar is 1 µm (E) Western blot of whole cell extracts from the indicated strains detecting TorA/TorΔE-GFP levels (α-GFP) with Ponceau stain to show relative protein loads. (F) Plot of SPB_f_/NE_f_ of TorA- and TorA-ΔE-GFP in individual cells from three biological replicates (n=32/replicate) with mean and SD. Kruskal Wallace ANOVA with *post hoc* Dunn’s test. ****=p<0.0001. (G, H) As in E, F.

### Outlook

In conclusion, we have employed the budding yeast model to interrogate the function of TorA. By uncovering an interaction between TorA and SPBs that can be altered by the introduction of the LAP1-LD, we have developed a visual *in vivo* assay to investigate the molecular determinants, and perhaps the function (in the future) of a TorA mediated ATP hydrolysis cycle. Perhaps most critically, our data point to the discovery of a likely substrate at the NE with the most obvious candidate being the SUN domain-containing Mps3. This interpretation is bolstered by our use of the yeast system in that there are no other conserved membrane proteins that anchor the SPB (as the mammalian centrosome does not span the NE), and budding yeast lack an obvious KASH domain partner (Mps2 is considered to be “KASH-like”(Friederichs *et al.*, 2011), but we did not identify any peptides to Mps2 in our affinity purifications). Indeed, the Mps3 SUN domain shares ∼30% sequence identity with its mammalian counterparts (Jaspersen *et al.*, 2006) and importantly, there are several additional studies that would support this conclusion (Nery *et al.*, 2008; Vander Heyden *et al.*, 2009; Jungwirth *et al.*, 2011; Atai *et al.*, 2012; Saunders *et al.*, 2017; Dominguez Gonzalez *et al.*, 2018). In the future, we look forward to the direct reconstitution of a SUN domain-TorA complex although it will be worthwhile to consider that the interaction between TorA and SUN-domain containing proteins need not be with the SUN domain per se as we also detect structural homology (with ∼99% confidence per Phyre2 (Kelley *et al.*, 2015)) between the recently identified auto-inhibitory domain in either Sun1 (Xu *et al.*, 2018) or Sun2 (Nie *et al.*, 2016) with the analogous region preceding the Mps3 SUN domain (Fig S2D-F).

Clearly, an additional priority going forward is to determine whether Pom152, and most critically the orthologous Nup210 directly interacts with TorA. The most exciting feature of this putative interaction, be it direct or indirect, is that it is specific for TorA and is disrupted by the ΔE mutation. These data are compelling as they lend support to the concept that defects in NPC biogenesis could arise in the absence of a putative TorA-Nup210 interaction and, in turn, contribute to the formation of the NE herniations that are the cellular hallmark of the dystonia disease phenotype. Perhaps most interestingly, our data hint at the existence of another co-factor that could be capable of stimulating TorA hydrolysis. While there is nothing obvious about the structure of the Pom152 lumenal domain, which consists of repetitive Ig motifs (Wozniak *et al.*, 1994; Upla *et al.*, 2017), that resembles the RecA fold of the LAP1 or LULL1 lumenal domains (Brown *et al.*, 2014; Sosa *et al.*, 2014), our data nonetheless raise the possibility that Pom152 could contribute to TorA-mediated ATP hydrolysis. Thus, we hope that this study facilitates direct testing of these hypotheses in mammalian systems and provides additional avenues to uncover the molecular basis of early onset DYT1 dystonia.

## ACKNOWLEDGEMENTS

We are grateful for invaluable discussions with Christian Schlieker and Megan King. This work was supported by the Dystonia Medical Research Foundation and the NIH: RO1 GM105672 to CPL. M.C. is also funded by the NIH 5F31HL134272 and 5T32GM007223 and K.W.B. by a National Science Foundation Graduate Research Fellowship DGE-1122492.

## Author contributions

M.C. and C.P.L. conceived of and designed all experiments, analyzed and interpreted all data and wrote the manuscript. K.W.B. aided in analysis of mass spec data. M.C. performed all experiments. S.B. and D.T. helped generate reagents.

## MATERIALS AND METHODS

### Plasmid generation

To generate pMC13/pMC25, pMC14/pMC26, and pMC22/pMC28, the coding sequences of TorA-GFP, TorA-ΔE-GFP, and TorA-EQ-GFP (Valastyan and Lindquist, 2011) (Addgene plasmid #32119, 32120, 32127) were subcloned into p406ADH1/p406GAL1 (gifts from Nicolas Buchler & Fred Cross; Addgene plasmid #15974, 17437) using *Xho*I and *Xba*I. The promoter and coding region of pMC13, pMC14, and pMC22 were then subcloned into pRS403 (ATCC) using *EagI* and *SacI* to generate pMC15, pMC16, and pMC24. The Gibson Assembly Mastermix (NEB) was used to insert the coding sequence for amino acids 328-583 of human LAP1 (LAP1-LD), which was PCR-amplified by Q5 DNA polymerase, into pDSL-NX (Dualsystems biotech, 2 *µ, TRP1*), which was used to generate NubG (N-terminus of Ubiquitin) fusions behind the control of the *CYC1* promoter, to generate pDT07. Subsequently, pMC12 was made by subcloning the NubG-HA-LAP1-LD coding sequence from pDT07 into p406GAL1 using *SpeI* and *SmaI*. pMC31, pMC33, and pMC35, encode the TorA-GD, LAP1-LD-R563A, and LAP1-LD-R563E point mutants, respectively, which were generated using Site Directed Mutagenesis with *Pfu Turbo* (Agilent Technologies).

### Yeast strain generation, growth and genetic analysis

All yeast strains are derived from a wt W303 genetic background and are listed in Supplemental Table 2. Gene knockouts and fluorescent protein/epitope-tagging of endogenous genes was performed using a PCR-based integration approach (Longtine *et al.*, 1998; Van Driessche *et al.*, 2005). To integrate TorA-GFP or LAP1-LD constructs into the genome at the *URA3* or *HIS3* loci, plasmids were digested with *BstBI* or *BmtI*, respectively, prior to transformation using standard methods (Sikorski and Hieter, 1989; Burke *et al.*, 2000). As indicated in Supplemental Table 2, some strains were generated by mating, subsequent sporulation and tetrad dissection using standard methods (Burke *et al.*, 2000). Yeast strains were grown at 30°C in YPA (1% yeast extract, 2% peptone, 0.025% adenine) with 2% dextrose (YPAD), 2% raffinose (YPAR) or 2% galactose (YPAG).

To test genetic interactions, relative growth of yeast strains was assessed by plating 10-fold serial dilutions of overnight cultures onto YPD or YPG plates. Plates were incubated at RT, 30ºC, 33ºC or 37 ºC and imaged at times indicated in figure legends.

### Preparation of whole-cell protein extracts and Western Blotting

To generate whole cell protein extracts for Western blotting, ∼2 OD_600_ of cells were collected by centrifugation, washed once with 1 mM EDTA, and lysed for 10 min on ice with 250 µL of 2M NaOH. Proteins were subsequently precipitated for 30 min after the addition of 250 µL 50% trichloroacetic acid. Samples were centrifuged at 18,000 g for 10 min at 4ºC to pellet precipitated proteins, washed once with 1 mL −20ºC acetone, resuspended in 40 µL 5% sodium dodecyl sulfate (SDS) followed by 40 µL of 2x SDS Laemmli sample buffer containing 100 mM dithiothreitol (DTT), and heat denatured for 5 min at 95 °C.

Protein samples were separated on a 4-20% gradient gel (BioRad) and transferred using the Mini Trans-Blot^®^ Cell (BioRad) at 100 V for 60 min onto 0.2 µm nitrocellulose (BioRad). Protein samples collected during affinity purification experiments (below) were separated on a 4-12% gel (NuPAGE) and transferred using Mini Blot Module (Invitrogen) at 25 V for 70 min onto 0.2 µm nitrocellulose (BioRad). Nitrocellulose was subsequently blocked with 5% skim milk in TBST for 1 h at RT, incubated with primary antibodies for 1 h at RT, extensively washed in TBST, incubated with HRP-conjugated secondary antibodies. After washing, proteins were detected by ECL. Antibodies used in this study are listed in Table 4.

### Affinity purification of TorA and TorA-ΔE-GFP

To identify the interactome of TorA-GFP and TorA-ΔE-GFP, 1 L cultures of W303 (No GFP control), MC99 (TorA-GFP), or MC108 (TorA-ΔE-GFP) were grown to an OD_600_ of ∼2, pelleted at 4000 g for 15 min at 4ºC. Cell pellets were transferred to a 50 mL conical tube, washed once with dH_2_O, weighed, and suspended in 100 µL freezing buffer (20 mM HEPES pH 7.5, 1.2% Polyvinylpyrrolidone, protease inhibitor cocktail (1:200)) per gram of cell pellet to make a slurry. A 20 gauge needle was used to puncture a hole into the bottom of a 50 mL conical tube and a plunger was used to force the slurry, drop-wise, through the hole into a second 50 mL conical tube filled with liquid nitrogen. Frozen cell pellets were cryomilled with a ball mill (Retsch) to generate a frozen powder of cell lysate (Alber *et al.*, 2007). For each affinity purification, 800 µL of extraction buffer (100 mM HEPES pH 7.5, 300 mM KCl, 1mM EDTA, 100 mM MgOAc, 1% triton X-100, 10 mM 2-mercaptoethanol was added to 200 mg of lysate powder. The sample was vortexed to resuspend prior to centrifugation at 14,000 rpm at 4ºC for 10 min to clear the lysate. The supernatant was added to 10 µL GFP-Trap^®^ magnetic-agarose beads (Chromotek) previously equilibrated in 1 mL lysis buffer. After a 1 hr incubation at 4ºC, the unbound fraction was collected, and the beads were washed 5x in 1 mL extraction buffer. To elute bound proteins, beads were resuspended in 20 µL lithium dodecyl sulfate sample buffer (NuPAGE), denatured at 70ºC for 10 min, centrifuged at 18,000 g for 1 min prior to physical separation of beads from the elution using a magnet. The elution was transferred to a fresh microcentrifuge tube, reduced by addition of 1 µL 1 M DTT (final concentration of 50 mM DTT), and heated again for 10 min at 70ºC. Bound proteins were identified by MS or Western Blotting.

### Mass Spectrometry

Eluted proteins were loaded onto a 4-12% gel (NuPAGE) until the dye front entered the gel. Proteins were visualized using Imperial Protein Stain (Thermo Scientific). After washing with dH_2_O, a clean razor blade was used to excise the protein band (∼10 µg). The gel piece was transferred to a clean 1.5 mL tube, and an in-gel tryptic digest was performed. Peptides were separated on a Waters nanoACQUITY ultra high pressure liquid chromatograph (UPLC) prior to detection on either a Waters/Micromass AB QSTAR Elite (two replicates for wt cells (no GFP) and those expressing TA-GFP; one replicate for cells expressing TorA-ΔE-GFP) or a ThermoScientific LTQ-Orbitrap XL Fusion (third replicate for wt (no GFP) and TA-GFP samples and second replicate for TorA-ΔE-GFP) mass spectrometer.

Mascot was used to analyze all MS/MS data (Matrix Science, London, UK; version 2.6.0). Mascot searched the SwissProt_2017_01 database (selected for *Saccharomyces cerevisiae*, unknown version, 7904 entries) assuming strict trypsin enzyme digestion, a fragment ion mass tolerance of 0.020 Da and allowed oxidation of methionine and propionamide of cysteine.

To validate MS/MS based peptide and protein identifications, Scaffold (version Scaffold_4.8.7, Proteome Software Inc., Portland, OR), a program that incorporates the Protein Prophet algorithm (Nesvizhskii *et al.*, 2003), was used with the following parameters: peptide identifications needed at least 95.0% probability by the Scaffold Local FDR algorithm; protein identifications required at least 22.0% probability to achieve an FDR less than 2.0% and at least 2 identified peptides; proteins that could not be distinguished by MS/MS analysis alone due to similar peptides were grouped to satisfy the principles of parsimony.

Scaffold (version Scaffold_4.8.7, Proteome Software Inc., Portland, OR) was used to generate normalized spectral counts for each replicate. To identify proteins at least 2.5-fold enriched in TorA-GFP or TorA-ΔE-GFP relative to the no GFP control, normalized peptide counts for individual proteins were averaged; TorA/ΔE-GFP average for each protein was divided by the corresponding average no GFP control. Proteins present in TorA/ΔE-GFP but absent in the no GFP control were also considered to be 2.5-fold enriched.

### Structure Prediction

To identify domain in Mps3 that are structurally similar to other SUN domain proteins, the region between the transmembrane domain and established SUN domain (Jaspersen *et al.*, 2006) (amino acids 171-458) were threaded into Phyre 2.0 (Kelley *et al.*, 2015). PyMOL was used to super-impose the predicted structure onto mouse Sun1 crystal structure (Xu *et al.*, 2018).

### Microscopy

A Deltavision widefield deconvolution microscope (Applied Precision/GE Healthcare) with a 100x, 1.40 numerical aperture objective (Olympus), solid-state illumination, and a CoolSnap HQ^2^ CCD camera (Photometrics) was used to acquire all images. To prepare cells for imaging, they were gently pelleted by centrifugation, suspended in complete synthetic medium (CSM) and pipetted onto a glass slide. Z-stacks with a 0.25 µm step were acquired with the exception of cells expressing TA-EQ-GFP: TA-EQ-GFP signal was low, so only a single plane was imaged to prevent signal loss due to photobleaching.

### Image processing, analysis and statistics

All images presented were deconvolved via softWoRx (Applied Precision GE Healthcare). Subsequent processing was performed using Fiji/ImageJ (Schindelin *et al.*, 2012) and Photoshop (Adobe). Image analysis (detailed below) was performed on raw (i.e. non-deconvolved) images using Fiji/ImageJ.

To determine the relative accumulation of TorA-GFP at the SPB, the GFP fluorescence at the SPB (SPB_f_) was divided by the NE GFP signal (NE_f_). To calculate SBP_f_, a six-by-six pixel square was used to measure the mean GFP fluorescence at the GFP accumulation that co-localized or over-lapped with the SPB, as visualized with Spc42-mCherry. To obtain a NE_f_ value, a portion of the NE was outlined and the mean GFP fluorescence was measured. After subtracting mean background fluorescence, SPB_f_ was divided by NE_f_ for each cell. To simplify the analysis, only cells with a single SPB were analyzed. 32 cells per genotype per biological replicate were analyzed, and each experiment represents 3 independent biological replicates, totaling in 96 cells per genotype per experiment.

The SPB_f_/NE_f_ distribution in TorA-GFP expressing cells failed the D’Agostino and Pearson normality test; therefore, the non-parametric Kruskal Wallace One-Way ANOVA was used to compare multiple genotypes within an experiment simultaneously and a *post hoc* Dunn’s test was used to identify significant differences.

---

**Figure S1:**
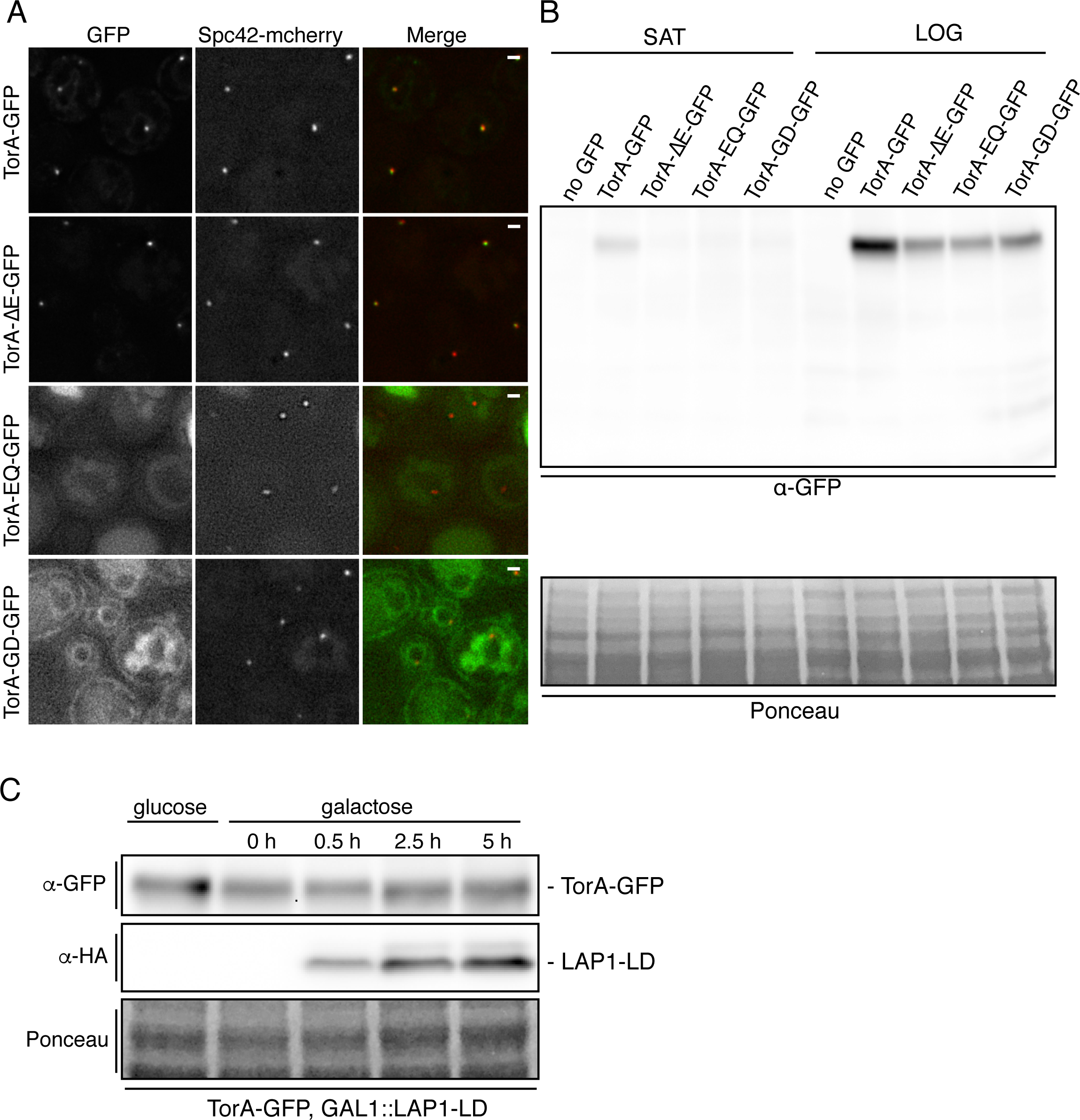
TorA-GFP and TorA-ΔE-GFP, but not TorA-GD-GFP accumulate at SPBs when produced at low levels. (A) Deconvolved fluorescent micrographs (green, red and merge) of strains expressing indicated TorA-GFP constructs and Spc42-mCherry after growth saturation. Bar is 1 µm. (B) Western blot of whole extracts from strains expressing indicated TorA-GFP constructs in saturated (SAT) and log phase cultures (LOG). Ponceau stain shows relative protein load. (C) Western blot of whole cell extracts from indicated strain under growth conditions that repress (glucose) or induce (galactose) the production of the LAP1-LD (detected by α-HA antibodies).

**Figure S2.**
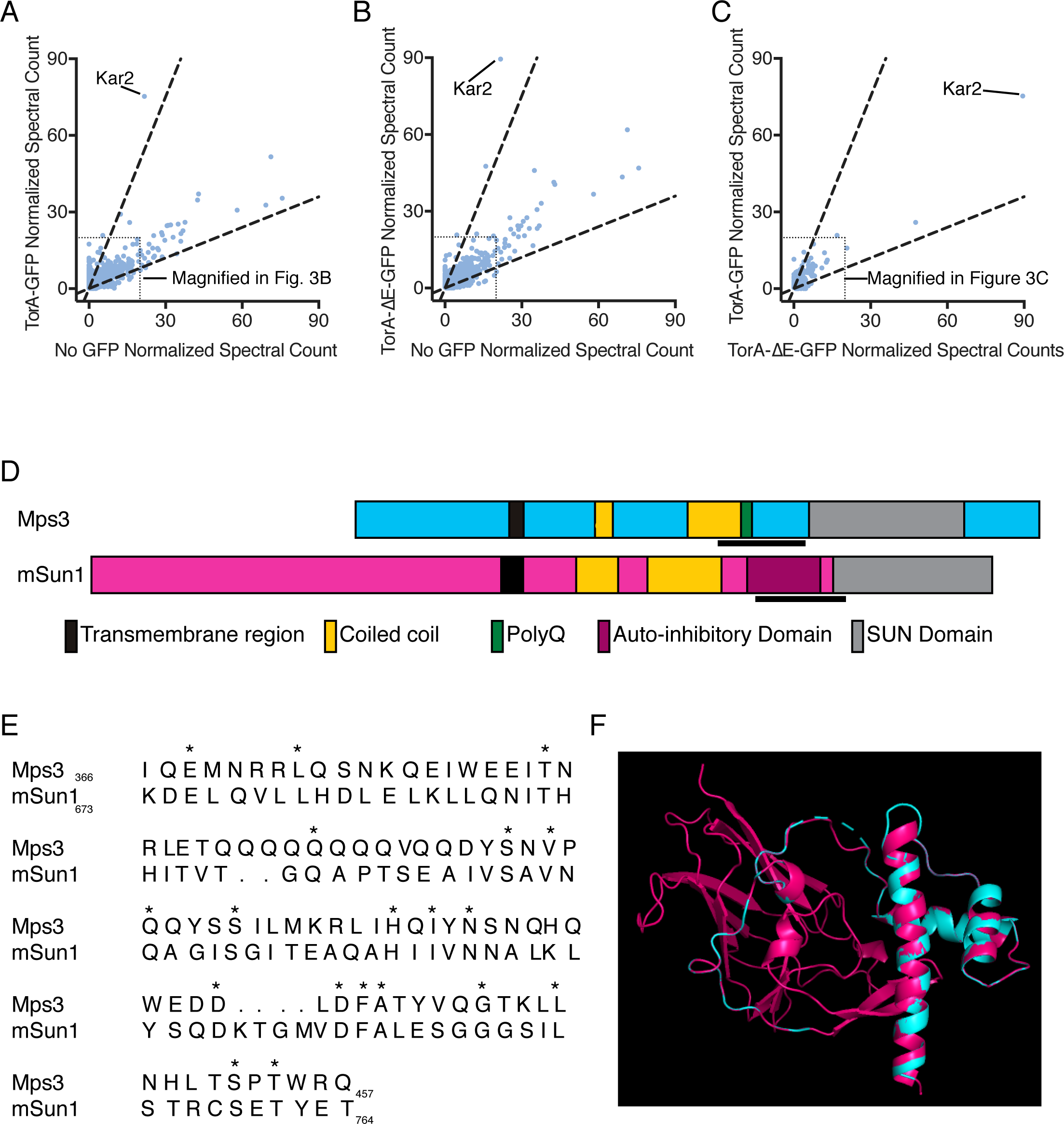
Identification of TorA-specific binding partners and analysis of Mps3 lumenal domain. (A-C) Plots of average normalized spectral counts from MS analysis of proteins derived from affinity purifications of TorA-GFP, TorA-ΔE-GFP and no GFP controls. Dashed lines demarcate 2.5-fold enrichment. Kar2 is indicated as the top hit. Boxed regions where the majority of peptides are found are magnified and presented in Figure 3. (D) Diagram comparing Mps3 (blue) to mouse (m) Sun1 (pink). The underlined region in mSUN1 is the autoinhibitory domain (amino acids 673-764). (E) Sequence alignment of the autoinhibitory domain of mSUN1 and the analogous region of Mps3 (amino acids 366-457). Asterisks denote identical amino acid residues. (F) Phyre2-generated structure prediction of the putative autoinhibitory region of Mps3 (blue) super-imposed onto the mouse autoinhibitory domain and SUN domain crystal structure (pink).

## REFERENCES

Adam, I., Jossé, L., and Tuite, M.F. (2017). Human TorsinA can function in the yeast cytosol as a molecular chaperone. Biochem J 474, 3439–3454.

Aitchison, J.D., Blobel, G., and Rout, M.P. (1995). Nup120p: a yeast nucleoporin required for NPC distribution and mRNA transport. J Cell Biol 131, 1659–1675.

Alber, F., Dokudovskaya, S., Veenhoff, L.M., Zhang, W., Kipper, J., Devos, D., Suprapto, A., Karni-Schmidt, O., Williams, R., Chait, B.T., Rout, M.P., and Sali, A. (2007). Determining the architectures of macromolecular assemblies. Nature 450, 683–694.

Atai, N.A., Ryan, S.D., Kothary, R., Breakefield, X.O., and Nery, F.C. (2012). Untethering the nuclear envelope and cytoskeleton: biologically distinct dystonias arising from a common cellular dysfunction. Int J Cell Biol 2012, 634214.

Brown, R.S., Zhao, C., Chase, A.R., Wang, J., and Schlieker, C. (2014). The mechanism of Torsin ATPase activation. Proc Natl Acad Sci U S A 111, E4822–4831.

Burke, D., Dawson, D., Stearns, T., and Cold Spring Harbor Laboratory. (2000). Methods in yeast genetics : a Cold Spring Harbor Laboratory course manual. Cold Spring Harbor Laboratory Press: Plainview, N.Y.

Chase, A.R., Laudermilch, E., Wang, J., Shigematsu, H., Yokoyama, T., and Schlieker, C. (2017). Dynamic functional assembly of the Torsin AAA+ ATPase and its modulation by LAP1. In: Mol Biol Cell, United States: 2017 by The American Society for Cell Biology.

Chen, J., Smoyer, C.J., Slaughter, B.D., Unruh, J.R., and Jaspersen, S.L. (2014). The SUN protein Mps3 controls Ndc1 distribution and function on the nuclear membrane. J Cell Biol 204, 523–539.

Chial, H.J., Rout, M.P., Giddings, T.H., and Winey, M. (1998). Saccharomyces cerevisiae Ndc1p is a shared component of nuclear pore complexes and spindle pole bodies. J Cell Biol 143, 1789–1800.

Crisp, M., Liu, Q., Roux, K., Rattner, J.B., Shanahan, C., Burke, B., Stahl, P.D., and Hodzic, D. (2006). Coupling of the nucleus and cytoplasm: role of the LINC complex. J Cell Biol 172, 41–53.

Demircioglu, F.E., Sosa, B.A., Ingram, J., Ploegh, H.L., and Schwartz, T.U. (2016). Structures of TorsinA and its disease-mutant complexed with an activator reveal the molecular basis for primary dystonia. Elife 5.

Dominguez Gonzalez, B., Billion, K., Rous, S., Pavie, B., Lange, C., and Goodchild, R. (2018). Excess LINC complexes impair brain morphogenesis in a mouse model of recessive TOR1A disease. Hum Mol Genet 27, 2154–2170.

Emtage, J.L., Bucci, M., Watkins, J.L., and Wente, S.R. (1997). Defining the essential functional regions of the nucleoporin Nup145p. J Cell Sci 110 (Pt 7), 911–925.

Friederichs, J.M., Ghosh, S., Smoyer, C.J., McCroskey, S., Miller, B.D., Weaver, K.J., Delventhal, K.M., Unruh, J., Slaughter, B.D., and Jaspersen, S.L. (2011). The SUN protein Mps3 is required for spindle pole body insertion into the nuclear membrane and nuclear envelope homeostasis. PLoS Genet 7, e1002365.

Goodchild, R.E., Buchwalter, A.L., Naismith, T.V., Holbrook, K., Billion, K., Dauer, W.T., Liang, C.C., Dear, M.L., and Hanson, P.I. (2015). Access of torsinA to the inner nuclear membrane is activity dependent and regulated in the endoplasmic reticulum. J Cell Sci 128, 2854–2865.

Goodchild, R.E., and Dauer, W.T. (2005). The AAA+ protein torsinA interacts with a conserved domain present in LAP1 and a novel ER protein. J Cell Biol 168, 855–862.

Goodchild, R.E., Kim, C.E., and Dauer, W.T. (2005). Loss of the dystonia-associated protein torsinA selectively disrupts the neuronal nuclear envelope. Neuron 48, 923–932.

Grillet, M., Dominguez Gonzalez, B., Sicart, A., Pöttler, M., Cascalho, A., Billion, K., Hernandez Diaz, S., Swerts, J., Naismith, T.V., Gounko, N.V., Verstreken, P., Hanson, P.I., and Goodchild, R.E. (2016). Torsins Are Essential Regulators of Cellular Lipid Metabolism. Dev Cell 38, 235–247.

Hanson, P.I., and Whiteheart, S.W. (2005). AAA+ proteins: have engine, will work. Nat Rev Mol Cell Biol 6, 519–529.

Hewett, J., Gonzalez-Agosti, C., Slater, D., Ziefer, P., Li, S., Bergeron, D., Jacoby, D.J., Ozelius, L.J., Ramesh, V., and Breakefield, X.O. (2000). Mutant torsinA, responsible for early-onset torsion dystonia, forms membrane inclusions in cultured neural cells. Hum Mol Genet 9, 1403–1413.

Jaspersen, S.L., and Ghosh, S. (2012). Nuclear envelope insertion of spindle pole bodies and nuclear pore complexes. Nucleus 3, 226–236.

Jaspersen, S.L., Giddings, T.H., and Winey, M. (2002). Mps3p is a novel component of the yeast spindle pole body that interacts with the yeast centrin homologue Cdc31p. J Cell Biol 159, 945–956.

Jaspersen, S.L., Martin, A.E., Glazko, G., Giddings, T.H., Jr., Morgan, G., Mushegian, A., and Winey, M. (2006). The Sad1-UNC-84 homology domain in Mps3 interacts with Mps2 to connect the spindle pole body with the nuclear envelope. J Cell Biol 174, 665–675.

Jokhi, V., Ashley, J., Nunnari, J., Noma, A., Ito, N., Wakabayashi-Ito, N., Moore, M.J., and Budnik, V. (2013). Torsin mediates primary envelopment of large ribonucleoprotein granules at the nuclear envelope. Cell Rep 3, 988–995.

Jungwirth, M., Dear, M.L., Brown, P., Holbrook, K., and Goodchild, R. (2010). Relative tissue expression of homologous torsinB correlates with the neuronal specific importance of DYT1 dystonia-associated torsinA. Hum Mol Genet 19, 888–900.

Jungwirth, M.T., Kumar, D., Jeong, D.Y., and Goodchild, R.E. (2011). The nuclear envelope localization of DYT1 dystonia torsinA-?E requires the SUN1 LINC complex component. BMC Cell Biol 12, 24.

Kelley, L.A., Mezulis, S., Yates, C.M., Wass, M.N., and Sternberg, M.J. (2015). The Phyre2 web portal for protein modeling, prediction and analysis. Nat Protoc 10, 845–858.

Kim, C.E., Perez, A., Perkins, G., Ellisman, M.H., and Dauer, W.T. (2010). A molecular mechanism underlying the neural-specific defect in torsinA mutant mice. Proc Natl Acad Sci U S A 107, 9861–9866.

Kustedjo, K., Bracey, M.H., and Cravatt, B.F. (2000). Torsin A and its torsion dystonia-associated mutant forms are lumenal glycoproteins that exhibit distinct subcellular localizations. J Biol Chem 275, 27933–27939.

Laudermilch, E., Tsai, P.L., Graham, M., Turner, E., Zhao, C., and Schlieker, C. (2016). Dissecting Torsin/cofactor function at the nuclear envelope: a genetic study. Mol Biol Cell 27, 3964–3971.

Liu, Q., Pante, N., Misteli, T., Elsagga, M., Crisp, M., Hodzic, D., Burke, B., and Roux, K.J. (2007). Functional association of Sun1 with nuclear pore complexes. J Cell Biol 178, 785–798.

Longtine, M.S., McKenzie, A., Demarini, D.J., Shah, N.G., Wach, A., Brachat, A., Philippsen, P., and Pringle, J.R. (1998). Additional modules for versatile and economical PCR-based gene deletion and modification in Saccharomyces cerevisiae. Yeast 14, 953–961.

Lu, W., Gotzmann, J., Sironi, L., Jaeger, V.M., Schneider, M., Lüke, Y., Uhlén, M., Szigyarto, C.A., Brachner, A., Ellenberg, J., Foisner, R., Noegel, A.A., and Karakesisoglou, I. (2008). Sun1 forms immobile macromolecular assemblies at the nuclear envelope. Biochim Biophys Acta 1783, 2415–2426.

Murphy, R., Watkins, J.L., and Wente, S.R. (1996). GLE2, a Saccharomyces cerevisiae homologue of the Schizosaccharomyces pombe export factor RAE1, is required for nuclear pore complex structure and function. Mol Biol Cell 7, 1921–1937.

Naismith, T.V., Dalal, S., and Hanson, P.I. (2009). Interaction of torsinA with its major binding partners is impaired by the dystonia-associated DeltaGAG deletion. J Biol Chem 284, 27866–27874.

Naismith, T.V., Heuser, J.E., Breakefield, X.O., and Hanson, P.I. (2004). TorsinA in the nuclear envelope. Proc Natl Acad Sci U S A 101, 7612–7617.

Nery, F.C., Zeng, J., Niland, B.P., Hewett, J., Farley, J., Irimia, D., Li, Y., Wiche, G., Sonnenberg, A., and Breakefield, X.O. (2008). TorsinA binds the KASH domain of nesprins and participates in linkage between nuclear envelope and cytoskeleton. J Cell Sci 121, 3476–3486.

Nesvizhskii, A.I., Keller, A., Kolker, E., and Aebersold, R. (2003). A statistical model for identifying proteins by tandem mass spectrometry. Anal Chem 75, 4646–4658.

Nie, S., Ke, H., Gao, F., Ren, J., Wang, M., Huo, L., Gong, W., and Feng, W. (2016). Coiled-Coil Domains of SUN Proteins as Intrinsic Dynamic Regulators. Structure 24, 80–91.

Ogura, T., Whiteheart, S.W., and Wilkinson, A.J. (2004). Conserved arginine residues implicated in ATP hydrolysis, nucleotide-sensing, and inter-subunit interactions in AAA and AAA+ ATPases. J Struct Biol 146, 106–112.

Onischenko, E., Tang, J.H., Andersen, K.R., Knockenhauer, K.E., Vallotton, P., Derrer, C.P., Kralt, A., Mugler, C.F., Chan, L.Y., Schwartz, T.U., and Weis, K. (2017). Natively Unfolded FG Repeats Stabilize the Structure of the Nuclear Pore Complex. Cell 171, 904-917.e919.

Ozelius, L.J., Hewett, J.W., Page, C.E., Bressman, S.B., Kramer, P.L., Shalish, C., de Leon, D., Brin, M.F., Raymond, D., Corey, D.P., Fahn, S., Risch, N.J., Buckler, A.J., Gusella, J.F., and Breakefield, X.O. (1997). The early-onset torsion dystonia gene (DYT1) encodes an ATP-binding protein. Nat Genet 17, 40–48.

Padmakumar, V.C., Libotte, T., Lu, W., Zaim, H., Abraham, S., Noegel, A.A., Gotzmann, J., Foisner, R., and Karakesisoglou, I. (2005). The inner nuclear membrane protein Sun1 mediates the anchorage of Nesprin-2 to the nuclear envelope. J Cell Sci 118, 3419–3430.

Rebelo, S., da Cruz E Silva, E.F., and da Cruz E Silva, O.A. (2015). Genetic mutations strengthen functional association of LAP1 with DYT1 dystonia and muscular dystrophy. Mutat Res Rev Mutat Res 766, 42–47.

Ryan, K.J., and Wente, S.R. (2002). Isolation and characterization of new Saccharomyces cerevisiae mutants perturbed in nuclear pore complex assembly. BMC Genet 3, 17.

Saunders, C.A., Harris, N.J., Willey, P.T., Woolums, B.M., Wang, Y., McQuown, A.J., Schoenhofen, A., Worman, H.J., Dauer, W.T., Gundersen, G.G., and Luxton, G.W. (2017). TorsinA controls TAN line assembly and the retrograde flow of dorsal perinuclear actin cables during rearward nuclear movement. J Cell Biol 216, 657–674.

Schindelin, J., Arganda-Carreras, I., Frise, E., Kaynig, V., Longair, M., Pietzsch, T., Preibisch, S., Rueden, C., Saalfeld, S., Schmid, B., Tinevez, J.Y., White, D.J., Hartenstein, V., Eliceiri, K., Tomancak, P., and Cardona, A. (2012). Fiji: an open-source platform for biological-image analysis. Nat Methods 9, 676–682.

Senior, A., and Gerace, L. (1988). Integral membrane proteins specific to the inner nuclear membrane and associated with the nuclear lamina. J Cell Biol 107, 2029–2036.

Sikorski, R.S., and Hieter, P. (1989). A system of shuttle vectors and yeast host strains designed for efficient manipulation of DNA in Saccharomyces cerevisiae. Genetics 122, 19–27.

Siniossoglou, S., Wimmer, C., Rieger, M., Doye, V., Tekotte, H., Weise, C., Emig, S., Segref, A., and Hurt, E.C. (1996). A novel complex of nucleoporins, which includes Sec13p and a Sec13p homolog, is essential for normal nuclear pores. Cell 84, 265–275.

Sosa, B.A., Demircioglu, F.E., Chen, J.Z., Ingram, J., Ploegh, H.L., and Schwartz, T.U. (2014). How lamina-associated polypeptide 1 (LAP1) activates Torsin. Elife 3, e03239.

Sosa, B.A., Rothballer, A., Kutay, U., and Schwartz, T.U. (2012). LINC complexes form by binding of three KASH peptides to domain interfaces of trimeric SUN proteins. Cell 149, 1035–1047.

Talamas, J.A., and Hetzer, M.W. (2011). POM121 and Sun1 play a role in early steps of interphase NPC assembly. J Cell Biol 194, 27–37.

Tanabe, L.M., Liang, C.C., and Dauer, W.T. (2016). Neuronal Nuclear Membrane Budding Occurs during a Developmental Window Modulated by Torsin Paralogs. Cell Rep 16, 3322–3333.

Thaller, D.J., and Patrick Lusk, C. (2018). Fantastic nuclear envelope herniations and where to find them. In: Biochem Soc Trans, vol. 46, England: 2018 The Author(s). Published by Portland Press Limited on behalf of the Biochemical Society., 877–889.

Upla, P., Kim, S.J., Sampathkumar, P., Dutta, K., Cahill, S.M., Chemmama, I.E., Williams, R., Bonanno, J.B., Rice, W.J., Stokes, D.L., Cowburn, D., Almo, S.C., Sali, A., Rout, M.P., and Fernandez-Martinez, J. (2017). Molecular Architecture of the Major Membrane Ring Component of the Nuclear Pore Complex. Structure 25, 434–445.

Valastyan, J.S., and Lindquist, S. (2011). TorsinA and the torsinA-interacting protein printor have no impact on endoplasmic reticulum stress or protein trafficking in yeast. PLoS One 6, e22744.

Van Driessche, B., Tafforeau, L., Hentges, P., Carr, A.M., and Vandenhaute, J. (2005). Additional vectors for PCR-based gene tagging in Saccharomyces cerevisiae and Schizosaccharomyces pombe using nourseothricin resistance. Yeast 22, 1061–1068.

Vander Heyden, A.B., Naismith, T.V., Snapp, E.L., Hodzic, D., and Hanson, P.I. (2009). LULL1 retargets TorsinA to the nuclear envelope revealing an activity that is impaired by the DYT1 dystonia mutation. Mol Biol Cell 20, 2661–2672.

VanGompel, M.J., Nguyen, K.C., Hall, D.H., Dauer, W.T., and Rose, L.S. (2015). A novel function for the Caenorhabditis elegans torsin OOC-5 in nucleoporin localization and nuclear import. Mol Biol Cell 26, 1752–1763.

Webster, B.M., Colombi, P., Jäger, J., and Lusk, C.P. (2014). Surveillance of nuclear pore complex assembly by ESCRT-III/Vps4. Cell 159, 388–401.

Webster, B.M., and Lusk, C.P. (2016). Border Safety: Quality Control at the Nuclear Envelope. Trends Cell Biol 26, 29–39.

Webster, B.M., Thaller, D.J., Jäger, J., Ochmann, S.E., Borah, S., and Lusk, C.P. (2016). Chm7 and Heh1 collaborate to link nuclear pore complex quality control with nuclear envelope sealing. EMBO J 35, 2447–2467.

Wente, S.R., and Blobel, G. (1993). A temperature-sensitive NUP116 null mutant forms a nuclear envelope seal over the yeast nuclear pore complex thereby blocking nucleocytoplasmic traffic. J Cell Biol 123, 275–284.

Wente, S.R., and Blobel, G. (1994). NUP145 encodes a novel yeast glycine-leucine-phenylalanine-glycine (GLFG) nucleoporin required for nuclear envelope structure. J Cell Biol 125, 955–969.

Winey, M., Hoyt, M.A., Chan, C., Goetsch, L., Botstein, D., and Byers, B. (1993). NDC1: a nuclear periphery component required for yeast spindle pole body duplication. J Cell Biol 122, 743–751.

Wozniak, R.W., Blobel, G., and Rout, M.P. (1994). POM152 is an integral protein of the pore membrane domain of the yeast nuclear envelope. J Cell Biol 125, 31–42.

Xiao, J., Bastian, R.W., Perlmutter, J.S., Racette, B.A., Tabbal, S.D., Karimi, M., Paniello, R.C., Blitzer, A., Batish, S.D., Wszolek, Z.K., Uitti, R.J., Hedera, P., Simon, D.K., Tarsy, D., Truong, D.D., Frei, K.P., Pfeiffer, R.F., Gong, S., Zhao, Y., and LeDoux, M.S. (2009). High-throughput mutational analysis of TOR1A in primary dystonia. BMC Med Genet 10, 24.

Xu, Y., Li, W., Ke, H., and Feng, W. (2018). Structural conservation of the autoinhibitory domain in SUN proteins. Biochem Biophys Res Commun 496, 1337–1343.

Yewdell, W.T., Colombi, P., Makhnevych, T., and Lusk, C.P. (2011). Lumenal interactions in nuclear pore complex assembly and stability. Mol Biol Cell 22, 1375–1388.

Zabel, U., Doye, V., Tekotte, H., Wepf, R., Grandi, P., and Hurt, E.C. (1996). Nic96p is required for nuclear pore formation and functionally interacts with a novel nucleoporin, Nup188p. J Cell Biol 133, 1141–1152.

Zacchi, L.F., Dittmar, J.C., Mihalevic, M.J., Shewan, A.M., Schulz, B.L., Brodsky, J.L., and Bernstein, K.A. (2017). Early-onset torsion dystonia: a novel high-throughput yeast genetic screen for factors modifying protein levels of torsinA?E. Dis Model Mech 10, 1129–1140.

Zacchi, L.F., Wu, H.C., Bell, S.L., Millen, L., Paton, A.W., Paton, J.C., Thomas, P.J., Zolkiewski, M., and Brodsky, J.L. (2014). The BiP molecular chaperone plays multiple roles during the biogenesis of torsinA, an AAA+ ATPase associated with the neurological disease early-onset torsion dystonia. J Biol Chem 289, 12727–12747.

Zhang, W., Neuner, A., Rüthnick, D., Sachsenheimer, T., Lüchtenborg, C., Brügger, B., and Schiebel, E. (2018). Brr6 and Brl1 locate to nuclear pore complex assembly sites to promote their biogenesis. J Cell Biol 217, 877–894.

Zhao, C., Brown, R.S., Chase, A.R., Eisele, M.R., and Schlieker, C. (2013). Regulation of Torsin ATPases by LAP1 and LULL1. Proc Natl Acad Sci U S A 110, E1545–1554.

Zhu, L., Millen, L., Mendoza, J.L., and Thomas, P.J. (2010). A unique redox-sensing sensor II motif in TorsinA plays a critical role in nucleotide and partner binding. J Biol Chem 285, 37271–37280.

Zirn, B., Grundmann, K., Huppke, P., Puthenparampil, J., Wolburg, H., Riess, O., and Müller, U. (2008). Novel TOR1A mutation p.Arg288Gln in early-onset dystonia (DYT1). J Neurol Neurosurg Psychiatry 79, 1327–1330.

